# BETA-HYDROXYBUTYRATE COUNTERACTS THE DELETERIOUS EFFECTS OF A SATURATED HIGH-FAT DIET ON SYNAPTIC AMPA RECEPTORS AND COGNITIVE PERFORMANCE

**DOI:** 10.1101/2024.01.23.576931

**Authors:** Rocío Rojas, Christian Griñán-Ferré, Aida Castellanos, Ernesto Griego, Marc Martínez, Juan de Dios Navarro-López, Lydia Jiménez-Díaz, José Rodríguez-Álvarez, David Soto del Cerro, Pablo E Castillo, Mercè Pallàs, Rut Fadó, Núria Casals

**Author notes:** Competing interest: The authors declare that no competing interests exist.

## Abstract

The ketogenic diet, characterized by high fat and low carbohydrates, has gained popularity not only as a strategy for managing body weight but also for its efficacy in delaying cognitive decline associated with neurodegenerative diseases and the aging process. Since this dietary approach stimulates the liver’s production of ketone bodies, primarily β-hydroxybutyrate (BHB), which serves as an alternative energy source for neurons, we investigated whether BHB could mitigate impaired AMPA receptor trafficking, synaptic dysfunction, and cognitive decline induced by metabolic challenges such as saturated fatty acids. Here, we observe that, in cultured primary cortical neurons, exposure to palmitic acid (200μM) decreased surface levels of glutamate GluA1-containing AMPA receptors, whereas unsaturated fatty acids, such as oleic acid and ω-3 docosahexaenoic acid (200μM), and BHB (5mM) increased them. Furthermore, BHB countered the adverse effects of palmitic acid on synaptic GluA1 levels in hippocampal neurons, as well as excitability and plasticity in hippocampal slices. Additionally, daily intragastric administration of BHB (100 mg/kg/day) for two months reversed cognitive impairment induced by a saturated high-fat diet (49% of calories from fat) in a mouse experimental model of obesity. In summary, our findings underscore the significant impact of fatty acids and ketone bodies on AMPA receptors abundance, synaptic function and neuroplasticity, shedding light on the potential use of BHB to delay cognitive impairments associated with metabolic diseases.

## Introduction

The escalating prevalence of cognitive impairment and dementia, projected to reach 150 million cases by 2050, represents a severe global public health concern (World Health Organization, n.d.). While increased life expectancy contributes to this outcome, attention has shifted to the impact of nutrition on brain health (Dye et al., 2017). Regular consumption of unhealthy saturated fatty acids (SFA), solid fats primarily derived from animals, and trans fats, liquid oils solidifying after food processing, has been associated with impaired memory performance, particularly in both young and old populations (Baym et al., 2014; Golomb & Bui, 2015; Okereke et al., 2012).

During cognitive tasks, neuroplasticity drives the formation and adjustment of synaptic connections, particularly within hippocampal glutamatergic circuits (Hanley, 2014). This fact involves precise control over the α-amino-3-hydroxy-5-methyl-4-isoxazolepropionic acid (AMPA)-type glutamate receptors (AMPARs) in the postsynaptic density (PSD). AMPARs are tetrameric assemblies of GluA1-4 subunits, being the GluA1/GluA2 heterotetramers, that are Ca^2+^-impermeable, the most common arrangement in mature neurons under resting conditions. Upon high neuronal activity, Ca^2+^-permeable GluA1 homomeric receptors are transiently incorporated at the plasma membrane (PM). This controlled GluA1-containing AMPARs inclusion supports synaptic potentiation during neuroplasticity, while a dysregulation of AMPAR subunit expression or trafficking imbalance can impact synaptic function, influencing cognitive processes (Jurado, 2018).

The impact of various fats on cognitive function is closely associated with alterations in synaptic function, plasticity, and modifications in synaptic AMPARs (Fadó et al., 2022). Diets rich in saturated fatty acids (SFAD) result in diminished spine density and narrowed PSD in the hippocampus (Granholm et al., 2008), leading to hyper-palmitoylation and hypo-phosphorylation of GluA1 (Spinelli et al., 2017). Treatment with saturated palmitic fatty acid (PA) reduces dendritic arborization and induces synaptic loss in hippocampal neurons, and both effects are successfully counteracted when co-treated with the ω-3 docosahexaenoic acid (DHA) (Loehfelm et al., 2020; McLean et al., 2019).

Recent work indicates that ketogenic diets, which are high in fats and low in carbohydrates, and ketone bodies produced by the liver during keto diet or fasting, mitigate cognitive decline associated with aging and neurodegeneration (Vinciguerra et al., 2020). Additionally, ketogenic formulas prevent memory deficits induced by hypoglycemia in type 1 diabetic patients (Page et al., 2009), while the administration of β-hydroxybutyrate (BHB), the primary KB, improves working memory in type 2 patients (Jensen et al., 2020). However, little is known about whether ketone bodies are capable of counteracting the pernicious effects of saturated fats on cognition and the implicated molecular mechanisms. Here, we explored the influence of BHB on AMPAR trafficking, synaptic plasticity, and cognition. Our results indicate that BHB is able to prevent PA-mediated surface withdrawal of GluA1 in cultured neurons and PA-mediated decrease in excitability and plasticity in hippocampal slices. Moreover, In SFAD-fed mice, daily intragastrical administration of BHB prevented diet-induced cognitive impairment, highlighting the potential of this ketone body in mitigating the detrimental effects of unhealthy nutrition on brain function.

## Results

### Effect of fatty acid and BHB on GluA1 surface levels in cortical neurons

Given the pivotal role of GluA1 in shaping synaptic plasticity, we explored the influence of saturated (PA) and the major ketone body, BHB, on GluA1 surface levels in vitro. Furthermore, the effects of unsaturated fatty acids, DHA and oleic acid (OA), were also analyzed. For this purpose, mouse primary cortical neurons cultured along 13-14 days *in vitro* (DIV) were treated with either PA (200 µM), OA (200 µM), DHA (200 µM), BHB (5 mM), or vehicle (0.6% BSA) in two distinct regimens: a brief exposure lasting 2 hours and a prolonged exposure spanning 24 hours. Then, we analyzed GluA1 levels on the PM by immunocytochemistry (IC) in nonpermeabilized cells using an antibody against its N-terminal extracellular domain.

Exposure of cortical cultures to PA reduced GluA1 levels at the PM (Figure 1A and Suppl. Figure 1), even with a short exposure. In contrast, OA or DHA treatment increased surface GluA1 (Figure 1B, C), with more pronounced effects with a longer exposure. Interestingly, BHB markedly enhanced GluA1 expression after both the 2 and the 24 hours of exposition (Figure 1D).

**Figure 1.**
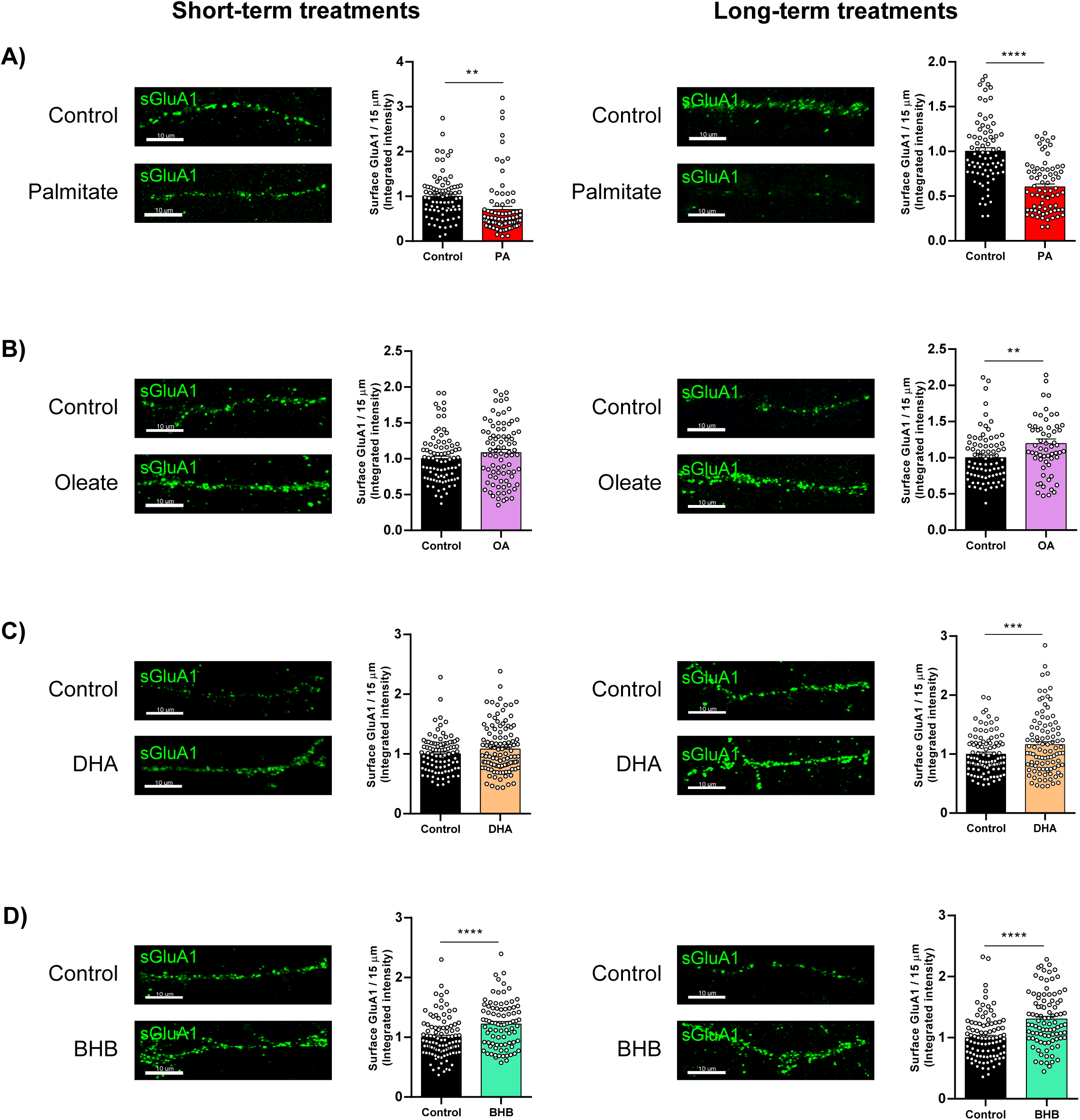
Nutrient’s effects on surface GluA1 levels in primary cortical neurons. Cortical neurons at 13-14 DIV were submitted to PA (200 µM; **A**), OA (200 µM; **B**), DHA (200 µM; **C**), or BHB (5mM; **D**) treatment, respectively. An equivalent amount of BSA solution was added to control cells when the effects of PA, OA, or DHA were analyzed or water, in the case of BHB. 2 and 24 hours later, neurons were fixed and processed by IC using an anti-GluA1 antibody against the N-terminal domain (extracellular; green stain). Representative images of dendrites are shown (see Supplemental Figure 1 for complete images). In each condition, we analyzed 45 neurons and 2 dendrites per neuron from three independent experiments performed by biological duplicates. The results in the graphs are the integrated intensity of surface GluA1 normalized by the control given as the mean ± SEM. (**A**) Left, control 2h (BSA; 1.00 ± 0.06, n = 79), and PA 2h (0.70 ± 0.07, n = 82); right, control 24h (BSA; 1.00 ± 0.04, n = 77), and PA 24h (0.60 ± 0.03, n = 72). (**B**) Left, control 2h (BSA; 1.00 ± 0.04, n = 84), and OA 2h (1.08 ± 0.04, n = 86); left, control 24h (BSA; 1.00 ± 0.04, n = 80), and OA 24h (1.19 ± 0.06, n = 60). (**C**) Left, control 2h (BSA; 1.00 ± 0.03, n = 93), and DHA 2h (1.08 ± 0.04, n = 114); right, control 24h (BSA; 1.00 ± 0.03, n = 100), and DHA 24h (1.16 ± 0.05, n = 99). (**D**) Left, control 2h (1.00 ± 0.04, n = 98), and BHB 2h (1.22 ± 0.04, n = 86); right, control 24h (1.00 ± 0.04, n = 95), and BHB 24h (1.30 ± 0.05, n = 93). **, P < 0.05; ***, P < 0.001; ****, P < 0.0001. Student t-test. Scale bar = 10 µm.

As an alternative experimental model, we also examined the effects of these nutrients in human-derived SH-SY5Y neuroblastoma cells. This cell line can be differentiated into a neuron-like phenotype upon exposition of retinoic acid, which promotes the growth of neurites. In this context, the treatment of differentiated SH-SY5Y with PA, OA, DHA, or BHB for 24 hours was able to recapitulate the same effects observed in cultured cortical neurons (Suppl. Figure 2). Therefore, these neuron-like cells are a suitable alternative *in vitro* model for studying the impact of these nutrients, avoiding the use of animals.

Altogether, these results confirm a distinct impact of PA, OA, DHA, and BHB on GluA1 surface levels. In addition, more potent effects were elicited with a long-term treatment than a shorter one.

### Synaptic transmission is reduced in palmitate-treated cortical neurons

To explore nutrient’s effects on synaptic transmission, we measured AMPA-mediated miniature excitatory postsynaptic currents (mEPSCs) in primary cortical neurons after the exposure to PA and BHB. We chose these two nutrients since they exhibited the most significant regulation on surface GluA1. To isolate AMPAR-mEPSCs, D-(-)-2- amino-5-phosphonopentanoic acid and picrotoxin were added to the perfusion to block N-methyl-D-aspartate (NMDA) and gamma-aminobutyric acid (GABA_A_) receptors, respectively.

AMPAR-mediated mEPSCs amplitudes were reduced in cortical neurons treated with PA for 24 hours, with no observable changes in the case of BHB (Figure 2A, B, C). Cumulative amplitude histograms also revealed a clear shift in the amplitude distribution towards smaller values under PA treatment (Figure 2D), consistent with a reduction in postsynaptic AMPAR number. In contrast, mEPSCs frequency was not significantly affected (Figure 2E), suggesting the absence of presynaptic alterations in neurons treated with PA or BHB. Kinetic parameters such as rise time and τ-weighted were unaffected by these treatments (Figure 2F, G), proving that synaptic AMPAR subtype composition remained unchanged. Therefore, exposure to PA decreases the abundance of postsynaptic AMPARs without detectable alterations in their subunit composition. The fact that BHB does not affect mEPSCs amplitude and frequency suggests that the increase it causes in surface GluA1 (Figure 1D) might be restricted to the perisynaptic region.

**Figure 2.**
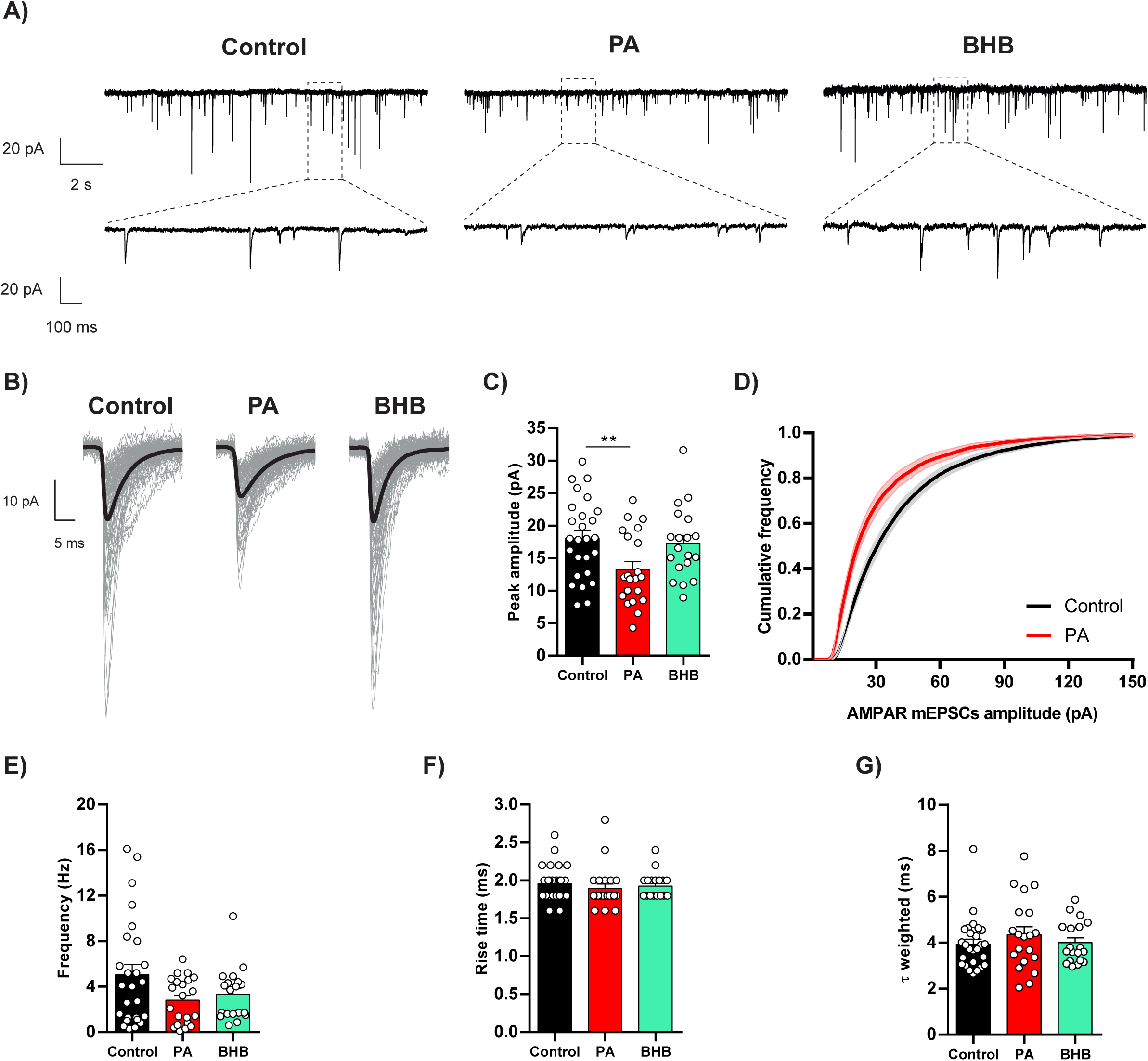
PA and BHB effects on mEPSC in cortical neurons. Cultured cortical neurons (14-15 DIV) were treated with PA (200µM) or BHB (5mM) for 24h. An equivalent amount of PA’s vehicle (BSA solution) was added to control neurons and BHB-treated ones to compare all the conditions properly. (**A**) Representative whole-cell recordings of AMPAR-mediated mEPSCs of 15 seconds duration. Inset magnifications correspond to 1.5 seconds. Membrane potential was held at -70 mV. (**B**) Examples of averaged mEPSCs (bold lines) from 11-, 7- and 5-minutes of time (control, PA, and BHB, respectively) of the recordings shown in A. Control neurons exhibited a mean mEPSCs amplitude of –18 pA (average of 2815 miniature events); PA-treated ones (central traces), –12.1 pA (1967 events); and BHB-treated ones, –18.38 pA (1098 events). One hundred random individual mEPSCs events are shown in grey for each group. (**C**) AMPAR mEPSCs amplitude was decreased in PA-treated cells compared with control neurons (-18.07 ± 1.20 pA, n=26; -13.28 ± 1.20 pA, n=21; and -17.25 ± 1.27 pA, n=19 for control, PA, and BHB-treated neurons respectively). (**D**) Cumulative probability distribution of mEPSCs amplitude showing a leftward shift in amplitude for events of PA-treated neurons. Continuous lines represent the average for control (black line; n=26 neurons; an average of 1104 events/neuron) and PA (red line; n=21; 768 events/neuron). Discontinuous lines denote the SEM. E) AMPAR mEPSCs frequency was not altered with PA nor with BHB (5.04 ± 0.92 Hz; 2.82 ± 0.44 Hz; and 3.32 ± 0.53 Hz; for control, PA, and BHB, respectively). (**F**) AMPAR mEPSCs rise time was not altered with PA nor with BHB (1.96 ± 0.05 ms; 1.90 ± 0.06 ms; and 1.93 ± 0.04 ms for control, PA, and BHB, respectively). (**G**) AMPAR mEPSCs τ-weighted was not altered either with PA nor with BHB (3.94 ± 0.21 ms; 4.35 ± 0.35 ms; and 4.00 ± 0.21 ms; for control, PA, and BHB, respectively). **, P < 0.01. Mann-Whitney test by comparing each condition with the control.

### BHB mitigates palmitate-mediated decrease in synaptic GluA1 levels

Given the increase of surface GluA1 levels under BHB treatment, we aimed to explore whether BHB could prevent PA-induced synaptic changes. To this end, we examined the effects of their co-treatment on GluA1 synaptic abundance by colocalizing this receptor with PSD95 as a postsynaptic density marker. Furthermore, we tested various concentrations of BHB (1 mM, 2 mM, and 5 mM), all falling within the high physiological range observed in mice (Nehlig, 2004). Here, mouse primary hippocampal cultures were employed to assess whether this neuronal type replicated the outcomes observed in cortical ones.

As expected, short-term exposition to PA reduced GluA1 surface levels in hippocampal neurons (Figure 3; upper graph; red bar) and the amount of synaptic GluA1 (Figure 3; lower graph;). Moreover, the treatment with PA led to a decrease in the levels of the synaptic marker PSD95 (Figure 3; middle graph; and Suppl. Figure 3). In contrast, BHB did not affect synaptic GluA1 abundance at any of the assessed concentrations (Figure 3; bellow graph; green bars), although surface GluA1 increased with 5 mM of BHB (Figure 3; upper graph), as previously observed in cortical neurons. These data align with the results on mEPSCs, where PA, but not BHB, induced alterations in AMPAR-mediated currents.

**Figure 3.**
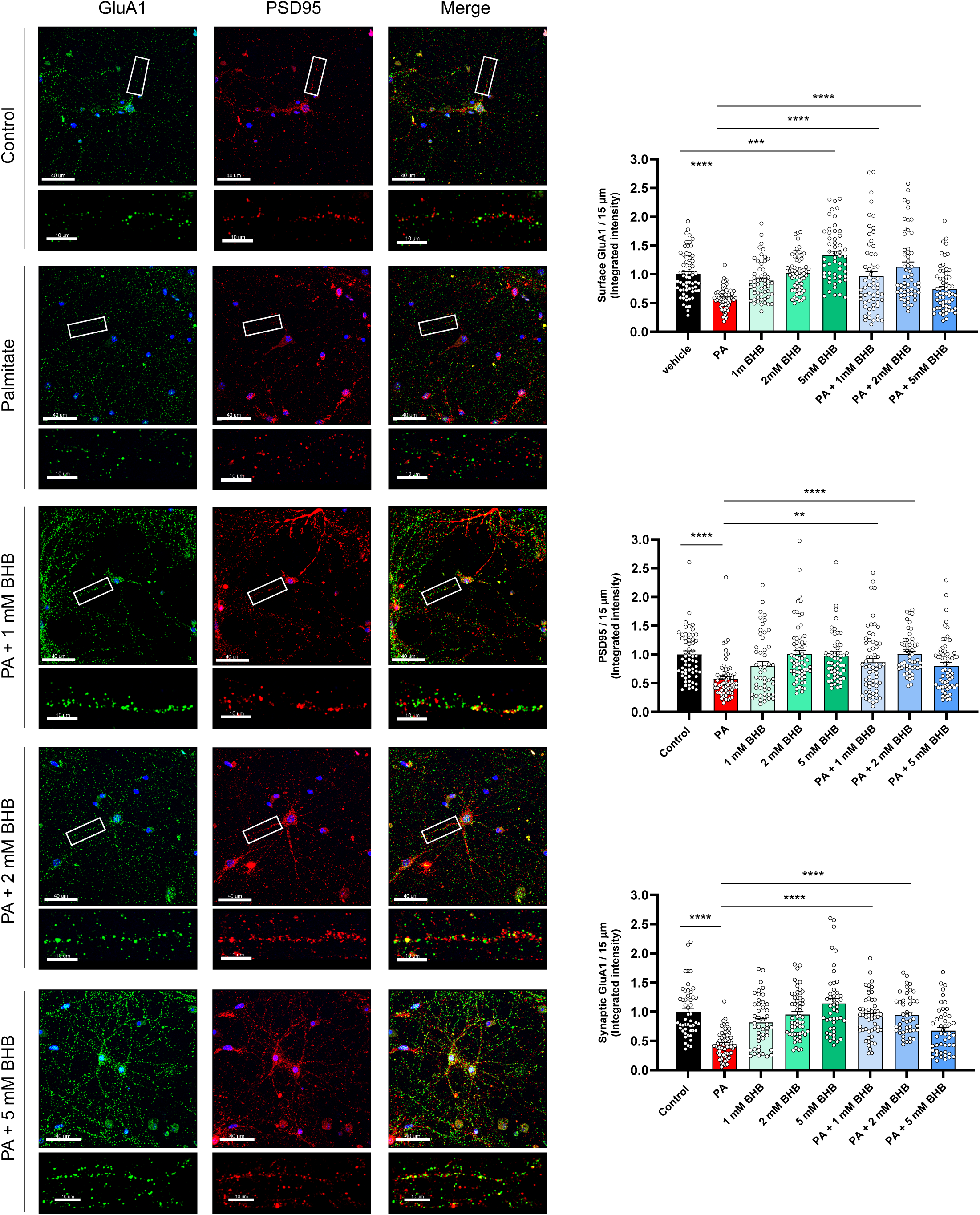
Short-term effects of PA and BHB on synaptic GluA1 levels in hippocampal neurons. Cultured hippocampal neurons (14-15 DIV) were treated with PA (200 µM) and/or BHB (1, 2 or 5 mM). An equivalent amount of vehicle (BSA solution) was added to control neurons and BHB-treated ones to properly compare all the conditions. 2 hours later, in non-permeabilized cells, GluA1 was detected by immunostaining. Then, cells were permeabilized, and a subsequent immunostaining was used to identify the synaptic marker PSD95. To determine synaptic GluA1 levels, the integrated intensity of GluA1 staining within the PSD95-defined ROI was quantified. Results in the graphs are given as mean ± SEM of 60 dendrites from two independent experiments performed by biological duplicates. Surface GluA1 (upper graph): Control (BSA; 1.00 ± 0.05, n=64), PA (0.59 ± 0.02, n=69), 1 mM BHB (0.89 ± 0.05, n=53), 2 mM BHB (1.02 ± 0.04, n=65), 5 mM BHB (1.34 ± 0.07, n=53), PA + 1 mM BHB (0.96 ± 0.08, n=60), PA + 2 mM BHB (1.31 ± 0.08, n=58), and PA + 5 mM BHB (0.75 ± 0.05, n=61). PSD95 (central graph): Control (BSA; 1.00 ± 0.06, n=61), PA (0.57 ± 0.04, n=63), 1 mM BHB (0.80 ± 0.08, n=51), 2 mM BHB (1.01 ± 0.06, n=68), 5 mM BHB (0.98 ± 0.07, n=56), PA + 1 mM BHB (0.86 ± 0.07, n=61), PA + 2 mM BHB (1.00 ± 0.05, n=53), and PA + 5 mM BHB (0.80 ± 0.06, n=63). Synaptic GluA1 (below graph): Control (BSA; 1.00 ± 0.06, n=49), PA (0.44 ± 0.03, n=58), 1 mM BHB (0.82 ± 0.06, n=45), 2 mM BHB (0.95 ± 0.05, n=59), 5 mM BHB (1.14 ± 0.08, n=46), PA + 1 mM BHB (0.92 ± 0.05, n=52), PA + 2 mM BHB (0.94 ± 0.05, n=45), and PA + 5 mM BHB (0.68 ± 0.06, n=47). Scale bar = 40 µm; scale bar of inset magnifications = 10 µm. **, P < 0.01; ***, P < 0.001; and ****, P < 0.0001. One-way ANOVA followed by the Bonferroni test.

To determine whether BHB could counteract the detrimental effects of PA on synaptic GluA1, hippocampal neurons were co-treated with both nutrients. We found that 1 mM and 2 mM of BHB effectively mitigated the adverse effects of PA on synaptic GluA1 and PSD95 levels, whereas 5 mM of BHB did not exhibit a significant beneficial impact (Figure3). Excessively high concentrations of BHB likely have indirect counterproductive effects at the synapse level. These findings suggest that the harmful effects of SFA on GluA1 synaptic abundance could be prevented through the co-administration of 1-2 mM BHB.

### BHB restores palmitate-induced impairment in hippocampal synaptic plasticity

To explore whether the beneficial effects of BHB following PA treatment extend to evoked synaptic transmission and synaptic plasticity, we performed extracellular field recordings of excitatory postsynaptic potentials (fEPSP) in the CA1 area of the hippocampus and whole-cell recordings from CA1 pyramidal cells in acute slices. Synaptic responses were elicited by stimulating Schaffer collateral (SC), and picrotoxin was included in the bath to block fast inhibitory synaptic transmission.

First, to test the effect of PA and BHB on basal synaptic transmission, input-output (I/O) curves were conducted by increasing stimulation intensity (from 10 to 90 V) in hippocampal slices following short-term exposition (2 hours) to PA (200 µM), BHB (2 mM), or a combination of both. Our results revealed a significant reduction in the slope of I/O curves under PA treatment compared with control slices, whereas BHB administration increased it (Figure 4A). Remarkably, PA and BHB co-administration demonstrated the BHB’s ability to counteract the effects induced by PA.

**Figure 4.**
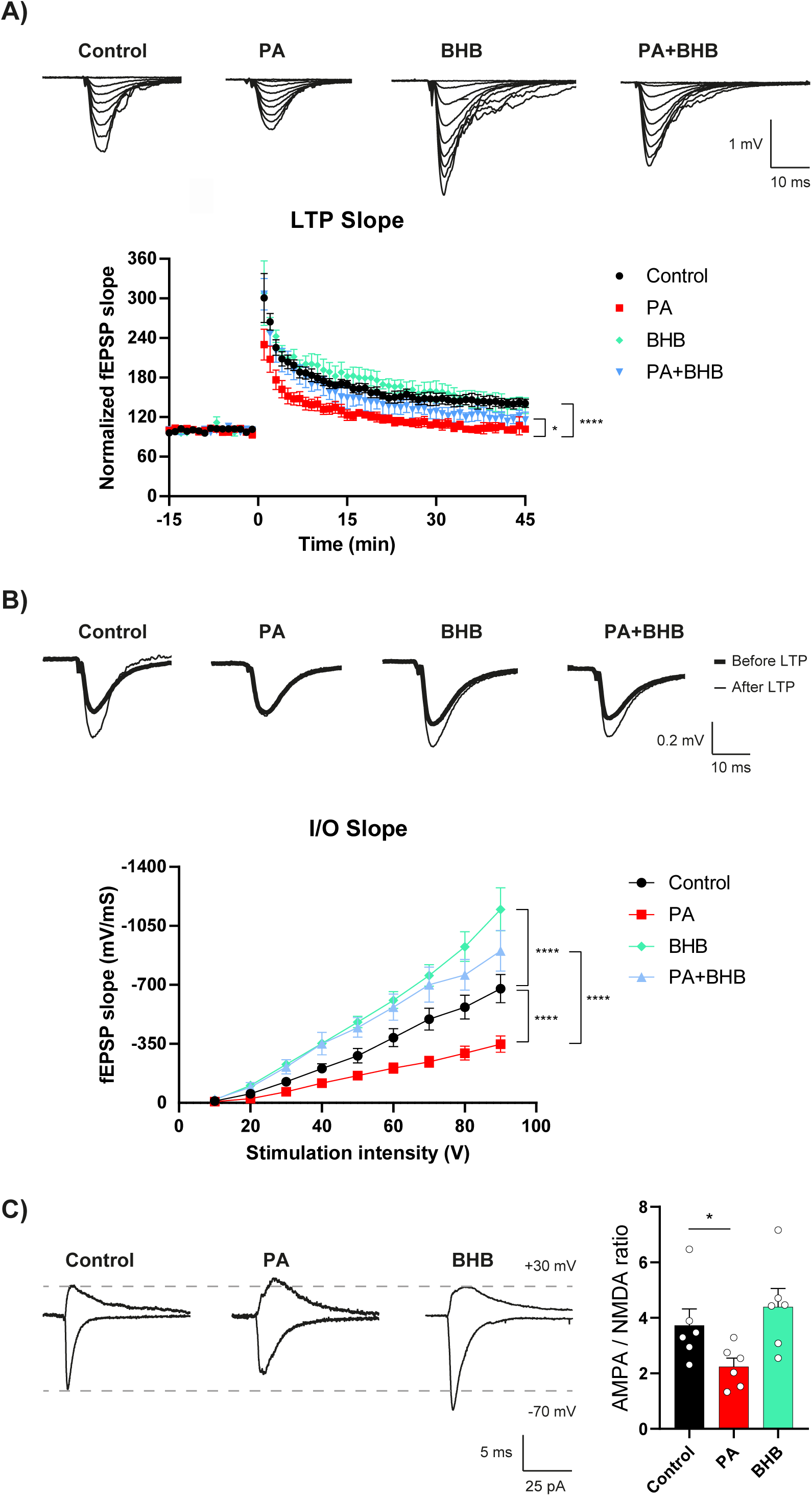
BHB reverses PA-induced impairment in electrophysiological I/O currents and LTP in hippocampal slices. (**A**) Superimposed fEPSP traces induced by a series of stimulating pulses with an increase of 10 mV applied to the SC fiber. Graph shows I/O curves plotted with the increased fEPSP slopes corresponding to the enhanced stimulating intensities. Results are the mean ± SD of n = 10-17 slices per condition taken from 4 male mice (P26 ± 5). Two-way ANOVA was followed by the Bonferroni. (**B**) Time course of normalized slope of AMPAR-mediated EPSCs from baseline and after induction of LTP. Results in the graphs are the mean ± SD of n = 3-4 slices per condition taken from 3-4 male mice (P27-34). One-way ANOVA was followed by the Bonferroni. Representative traces of the baseline (thin lines) and the last 5 minutes of the time course after LTP induction (thick lines). (**C**) Representative traces of AMPA and NMDA receptor-mediated postsynaptic currents were recorded at -70 and +30 mV, respectively. Results in the graphs showing the effect of the different treatments on the AMPA/NMDA ratio are represented as mean ± SEM, n=5-6 cells from 3-5 male mice (P26 ± 5) per condition. Control (3.73 ± 0.59, n=6), vehicle (4.16 ± 0.74, n=5), PA (2.25 ± 0.31, n=6), and BHB (4.40 ± 0.66, n=6). Student t-test by comparing each condition with the control. *, P < 0.05; **, P < 0.01; and ****, P < 0.0001.

We next analyzed the effect of PA and BHB on CA1 LTP. After a stable 15-minute baseline was established, long-term potentiation (LTP) was induced by high-frequency stimulation. BHB alone did not significantly affect the LTP magnitude (Figure 4B), however it was able to successfully mitigate the LTP reduction triggered by PA, when simultaneously administered. To test whether synaptic plasticity changes had a presynaptic origin, we measured the paired-pulse (PPR), a metric linked to the probability of neurotransmitter release. PPR was unaffected across all analyzed conditions (Suppl. Figure 4A, B, C), indicating that the effects of PA and BHB are likely postsynaptic.

Finally, we calculated the AMPA/NMDA ratio by comparing the amplitudes of EPSCs mediated by AMPA and NMDA receptors. PA significantly reduced the AMPA/NMDA ratio, whereas BHB induced a modest upward trend (Figure 4C). These results suggest that the PA-mediated effect in synaptic transmission is likely due to a reduction in AMPARs but not in NMDA receptors.

In summary, SFA and ketone bodies affected postsynaptic but not presynaptic function. In addition, BHB restored the PA-mediated impairment in synaptic transmission and plasticity. Consistent with our observations in cultured neurons, our findings using hippocampal slices support the notion that PA and BHB effects are mediated mostly through AMPARs.

### BHB counteracts the cognitive impairment mediated by a saturated fatty acid-rich diet in mice

To assess whether the beneficial effects of BHB extend to mitigating PA-induced cognitive impairment, we conducted *in vivo* experiments in mice. Specifically, we explored whether daily administration of BHB could counteract the adverse effects of a SFAD on memory task performance. To achieve this goal, 5 weeks-old mice were fed SFAD (constituting 49% of calories from SFA) or a chow diet (CD; 10% of calories from fat) for seven weeks. Based on previous studies by (Hu et al., 2018), we decided to perform an intragastrically administration of 100 mg/kg/day BHB in mice or an equal volume of vehicle (water) throughout this period. This dosage increased BHB levels within the hippocampus, reaching a maximum within 4 hours post-administration (Suppl. Figure 5). Body weight and food intake were monitored weekly. Finally, behavior tests were performed, and the animals were euthanized to obtain brain tissue, as illustrated in the experimental chronogram (Figure 5A).

**Figure 5.**
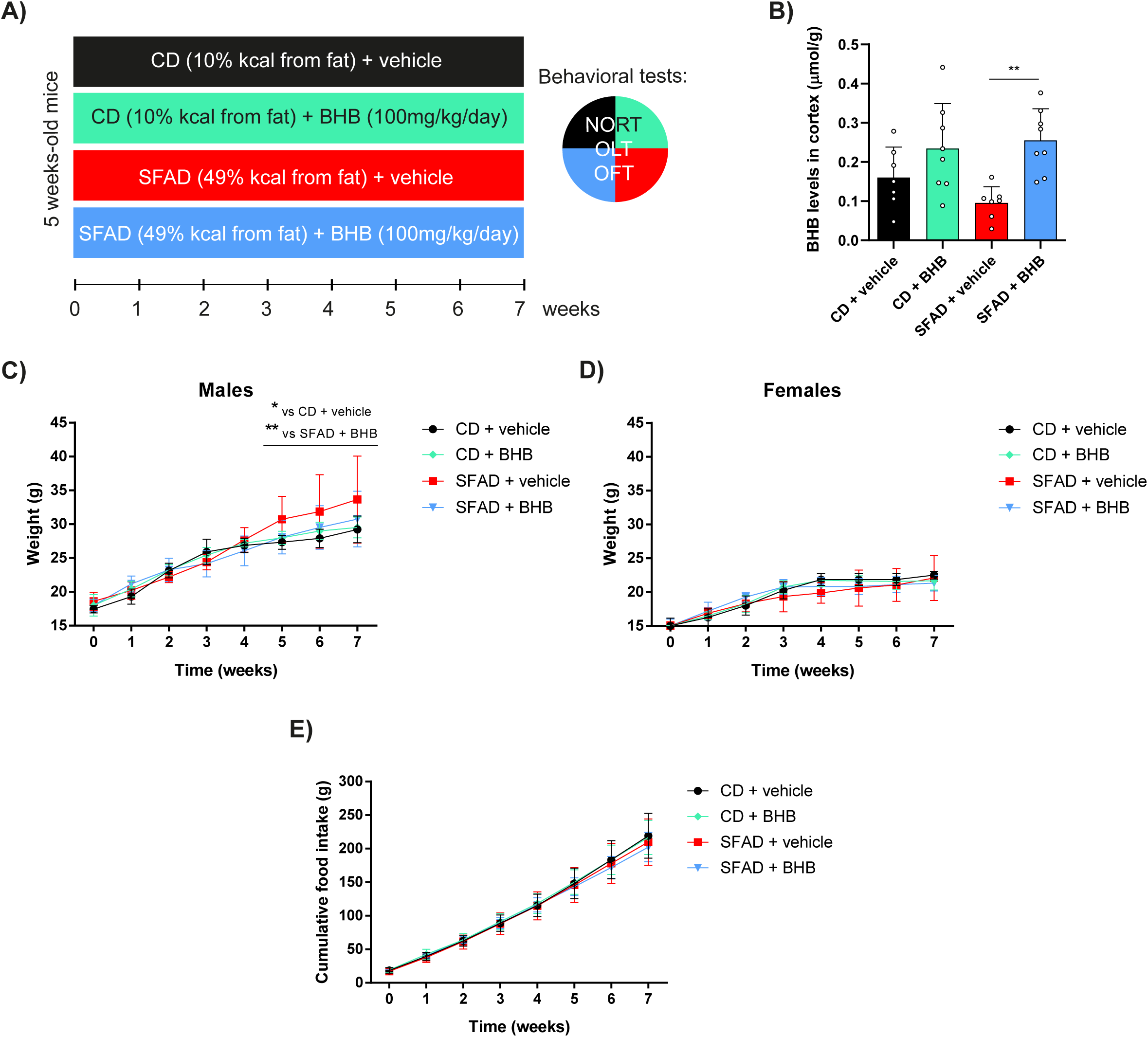
Effect of BHB administration in bodyweight gain mediated by SFAD. (**A**) Animals’ study design. 5-weeks old-mice were divided into 4 groups: CD (10% of kcal from FA) + vehicle (equal amount of water), CD + BHB administration (100mg/kg/day), SFAD (49% of kcal from SFA) + vehicle, and SFAD + BHB. Animals were maintained for 7 weeks under these feeding conditions and submitted to behavior test 3 days before the sacrifice to obtain blood and tissue samples. (**B**) The BHB levels in hippocampi. The results in the graphs are the mean ± SD. CD + vehicle (0.16 ± 0.08, n=7), CD + BHB (0.23 ± 0.12, n=8), SFAD + vehicle (0.10 ± 0.04, n=7), and SFAD + BHB (0.26 ± 0.08, n=8). (**C, D**) The animals were weighed weekly. Male and female weight is shown in C and D, respectively. Data are represented as mean ± SD from n=5-7/group. (**E**) The food was weighed weekly. Cumulative food intake is shown in g. Data are represented as mean ± SD from n=9-11/group. *, P < 0.05; and **, P < 0.01. One-way ANOVA was followed by the Bonferroni.

BHB levels were measured in the cortical region at the end of the experiment. As depicted in Figure 5B, the cortex exhibited elevated BHB levels in animals administered with BHB compared to those receiving the vehicle. These differences were significant in the high-fat diet-fed animals. However, the SFAD *per se* failed to raise brain BHB levels compared to a CD, contrasting with previous observations where high fat diet was associated with increased BHB brain levels (Jensen et al., 2020). This suggests that the particular response might depend on experimental conditions and diet-specific composition.

Throughout the experiment, the body weight revealed a tendency for males to exhibit higher weight when subjected to SFAD, although this difference only reached statistical significance from weeks 5 to 7 (Figure 5C, D). This effect was absent in females, a finding consistent with the well-documented hormone-dependent resistance of adult females to diet-induced obesity compared to males (Fadó et al., 2022). Notably, male mice supplemented with BHB did not display this trend of weight gain. Since the cumulative food intake was the same in all the groups (Figure 5E), results suggest that BHB prevents body weight gain triggered by SFAD in male mice.

To examine cognitive performance of mice, two behavior tests were conducted at the end of the experiment: novel object recognition test (NORT) and object location test (OLT). NORT included short-term memory (2 hours post-habituation) and long-term memory (24 hours later), both assessed using the discrimination index (DI), a metric reflecting the animal’s ability to recognize and remember a previously encountered object. While no significant differences were observed among CD-fed groups, mice exposed to SFAD exhibited substantial cognitive impairments completely prevented by concurrent BHB administration (Figure 6A, B). In the assessment of spatial learning evaluated through open field test (OLT), a remarkable difference was also found between CD and SFAD (Figure 6C), corroborating the detrimental effects of SFA on brain functions. Once more, BHB co-administration demonstrated its efficacy in counteracting SFAD harmful effects. There was no significant difference in total exploration time among the groups, and the same effects were observed in both females and males. In this context, the examination of locomotor activity revealed that, when administered with SFAD, BHB increased the distance traveled by mice compared to the other experimental groups (Figure 6D), suggesting heightened locomotor activity or exploration behavior and reduced anxiety, as demonstrated previously in animal models of stress (Yamanashi et al., 2020). By contrast, there were no differences in the number of rearings across all the groups (Figure 6E).

**Figure 6.**
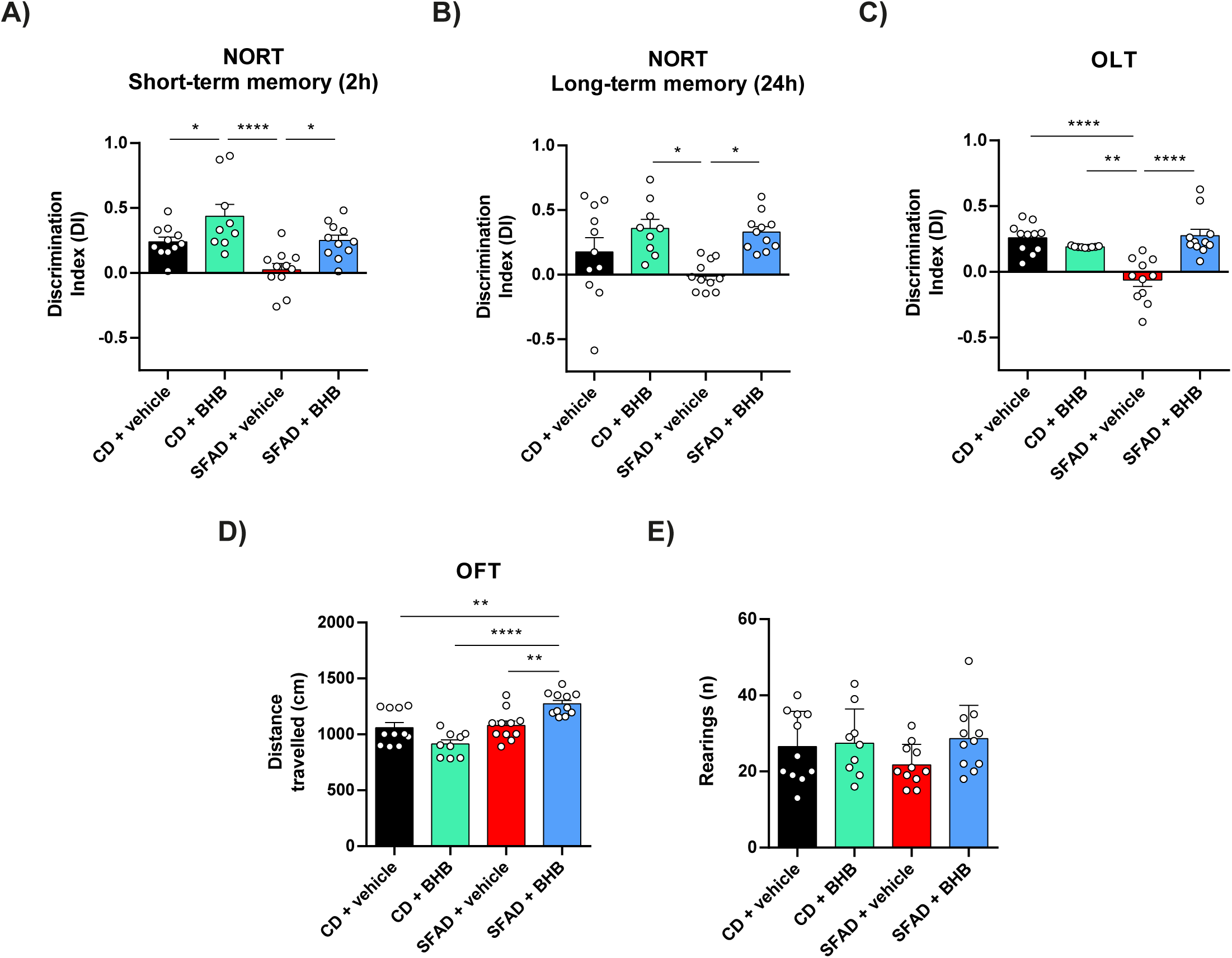
Effect of BHB administration in cognitive impairment mediated by SFAD. (**A, B, C**) NORT was performed 2h (**A**) and 24 hours (**B**) after the habituation phase, respectively. OLT was performed to evaluate spatial memory (**C**). Total exploration time was not significantly different among groups (data non-shown). The results of the DI in the graphs are the mean ± SD. NORT 2h: CD + vehicle (0.24 ± 0.12, n=11), CD + BHB (0.44 ± 0.28, n=9), SFAD + vehicle (0.02 ± 0.16, n=11), and SFAD + BHB (0.25 ± 0.14, n=11). NORT 24h: CD + vehicle (0.18 ± 0.37, n=11), CD + BHB (0.36 ± 0.21, n=9), SFAD + vehicle (-0.01 ± 0.12, n=11), and SFAD + BHB (0.33 ± 0.14, n=11). OLT: CD + vehicle (0.26 ± 0.03, n=11), CD + BHB (0.19 ± 0.002, n=9), SFAD + vehicle (-0.06 ± 0.05, n=11), and SFAD + BHB (0.26 ± 0.05, n=11). (**D, E**) Locomotor activity and the number of rearings were measured using OFT. Data are represented as mean ± SD. Locomotor activity: CD + vehicle (1059 ± 154, n=11), CD + BHB (914 ± 109, n=9), SFAD + vehicle (1079 ± 134, n=11), and SFAD + BHB (1274 ± 101, n=11). Rearings: CD + vehicle (26.5 ± 9.2, n=11), CD + BHB (27.4 ± 9.0, n=9), SFAD + vehicle (21.7 ± 5.4, n=11), and SFAD + BHB (28.6 ± 8.8, n=11). **, P < 0.01; and ****, P < 0.0001. One-way ANOVA followed by Bonferroni test.

Collectively, our findings reveal the beneficial role of BHB supplementation in counteracting the detrimental effects of a high-saturated-fat diet on cognitive abilities in mice.

## Discussion

There is a noteworthy surge in the popularity of high-fat, low-carbohydrate diets for body weight management and obesity and diabetes treatment (Basolo et al., 2022). These diets are widely reported to be effective in improving body composition parameters as well as glycemic and lipid profiles (Muscogiuri et al., 2021; Zhou et al., 2022). However, to the best of our knowledge, there is limited data on the beneficials effects of ketone bodies in reversing cognitive damage mediated by nutrients such as SFA. In the present study, we demonstrate that intragastric administration of BHB successfully counteracts the memory and learning impairments induced by SFAD in a mouse experimental model. We attribute these effects to the ability of BHB to prevent PA-induced adverse changes at the synaptic level.

Alterations in the expression of GluA1 or deregulations of its trafficking towards the PM are linked to impaired synaptic plasticity and cognition (Hanley, 2014; Jurado, 2018). Our study reveals that PA treatment significantly decreases surface GluA1 levels, while OA, DHA, and BHB increase them in hippocampal and cortical cultured neurons.

Our findings align with existing literature showing the beneficial effects of unsaturated fatty acids on cognition. Specifically, ω-3 supplementation in rodents enhances AMPAR levels, dendritic spine density, and LTP (Cansev et al., 2008; Kavraal et al., 2012; Lee et al., 2012), while ω-3 deficiency in pups reduces GluA1 in hippocampal synaptosomes and CA1-hippocampal LTP (Aryal et al., 2019). Moreover, surface GluA1 in hippocampal neurons is modulated by DHA (Ménard et al., 2009). In other studies, it successfully prevented the detrimental effects of PA in co-treated hippocampal and cortical neurons (Loehfelm et al., 2020; McLean et al., 2019). Therefore, we believe that DHA and OA, by enhancing the surface abundance of GluA1, might contribute to mitigate the negative impact of SFA on cognition. This is especially relevant in mixed diets, such as the Mediterranean diet, which is richer in polyunsaturated fatty acids (PUFAs)/ monounsaturated fatty acids (MUFAs) than in SFA, thus proving beneficial for cognition in the context of metabolic and neurodegenerative diseases (Canhada et al., 2018; Fadó et al., 2022; Kouvari et al., 2022; Vinciguerra et al., 2020).

Regarding PA, previous studies showed its ability to reduce AMPA sensitivity in cortical neurons, with no observable changes on NMDA sensitivity (Loehfelm et al., 2020), suggesting a specific effect of PA on AMPARs, in accordance with our results. Other investigations have examined the impact of PA in combination with insulin, revealing that the co-treatment caused GluA1 hyper-palmitoylation, concurrently downregulating its recruitment to the PM, and inhibiting AMPAR currents and their potentiation (Spinelli et al., 2017). Our study demonstrates that PA *per se* is sufficient to reduce both surface and synaptic levels of GluA1, resulting in a decrease in AMPARs-mediated currents and synaptic plasticity.

By contrast, the influence of BHB supplementation on surface or synaptic GluA1 levels has not been previously evaluated. Here, we demonstrate that BHB treatment, even with short exposures, can counteract the negative effects of PA on GluA1 surface and synaptic levels, AMPAR-mediated currents, and synaptic plasticity. It is important to notice that BHB alone has no effects on AMPAR synaptic abundance or plasticity. However, high doses of BHB increase surface GluA1 levels. These results suggest that the BHB-mediated increase in surface GluA1 might be confined to the perisynaptic zone—a subset of GluA1-containing AMPARs that could be swiftly and transiently mobilized to synapses under specific conditions, being involved in the reversibility of both synaptic and dendritic spine modifications (Yang et al., 2008, 2010).

Furthermore, PA and BHB effects on synaptic transmission and plasticity are mainly postsynaptic, as supported by the lack of alterations in the PPR and mEPSC frequency. Moreover, the reduction in the AMPA/NMDA ratio observed in response to PA indicates that NMDA receptors are likely not involved in PA-triggered synaptic changes. Our findings are consistent with previous studies where PA reduced LTP without affecting NMDA-mediated long-term depression (Contreras et al., 2017; Spinelli et al., 2017). Moreover, mEPSC data point to an unchanged subtype composition, suggesting that the trafficking of all AMPAR subunits is affected similarly by PA treatment. Interestingly, a high-fat diet induces both GluA1 and GluA2 palmitoylation (Spinelli et al., 2017), a post-translational modification that decreases the surface expression of both subunits (Hayashi et al., 2005), which is in supported by our results.

Noteworthy, other studies obtained similar results in a different context. BHB treatment effectively prevented the impartment of LTP mediated by several injuries in hippocampal slices (such as exposure to excitotoxic levels of NMDA, rotenone toxin or hydrogen peroxides, without inducing any alterations when administrated alone (Kimura et al., 2012; Maalouf & Rho, 2008; Youssef, 2015). Furthermore, in a mouse model of Alzheimer’s disease, BHB-mediated improvements in cognition were associated with increased LTP, while in young or healthy animals, a ketogenic diet unaffected or even decreased LTP (Blaise et al., 2015; Yin et al., 2016). Shortly after the induction of LTP, AMPARs are transported via exocytosis to perisynaptic sites, before their translocation to the synaptic zone for full LTP expression (Yang et al., 2008). BHB might increase AMPAR trafficking to the surface and facilitate a larger pool of GluA1 to be readily available in the perisynaptic area. In specific damaging conditions, such as a surplus of SFA, these receptors would then move rapidly laterally to the synapse, restoring normal synaptic function.

Exposure to obesogenic dietary components, like SFA, contributes to cognitive decline compared to isocaloric standard diets at any life stage (Fadó et al., 2022). As expected, in our study, the consumption of SFAD (49% of kcal from fat and 31% from carbohydrates) in mice induced memory and learning impairments in both sexes equally, although weigh gain was increased in males but not in female mice. This underscores that the cognitive negative impact may precede or surpass the adverse metabolic effects of SFAD. Similar results were obtained with a short-term administration of a high-fat, high-sucrose diet, which caused cognitive impairment in the absence of overt metabolic disturbances (Ramírez et al., 2022).

Despite the great popularity of the ketogenic diet as a strategy for managing body weight, evidence demonstrates that it is equally effective for weight loss as diets with higher carbohydrate content (Fernández-Verdejo et al., 2023). The consumption of ketone esters as supplements (often BHB linked to an ester molecule) has been associated with satiety and increased energy expenditure, particularly in the presence of high-carbohydrate availability, contributing to an enhancement in weight loss. Similarly, in our study, the administration of BHB exhibited a reduction in body weight in male mice fed with SFAD, which might be due to increased energy expenditure, as suggested by the heightened locomotor activity.

BHB *per ser* under a standard diet has an effect on short-term memory but cannot induce a sustained enhancement in spatial learning and long-term memory. These results align perfectly with a previous report where BHB supplementation improved short-term memory, but not the long-term, in healthy animals (Yin et al., 2016). Interestingly, we took a significant step beyond previous publications by demonstrating that BHB administration successfully prevented SFAD’s harmful effect on cognition. The oral administration of a single daily dose of BHB was sufficient to neutralize the negative effects of a high-fat diet on both short- and long-term memory, as well as exploratory interest, in adult mice.

Our findings point out that the cognitive benefits observed with ketogenic diets may be (at least partially) mediated by the ketone bodies produced during their consumption, which counteract the harmful effects of the SFA in the diet. Interestingly, ketogenic-based formulas used as supplements in diets more consistently enhance learning and memory performance than ketogenic diets, especially in older populations and Alzheimer’s disease patients (Fadó et al., 2022). Furthermore, our results provide a potential explanation for the cognitive benefits induced by intermittent fasting (Phillips, 2019). During fasting, the hypoglycemia response triggers the release of SFA from adipose tissue into the bloodstream, while the liver produces ketone bodies. We propose that, in such situations, the elevated levels of circulating ketone bodies not only serve as the primary energy source for neurons but also mitigate the negative impact of SFA on synaptic function and cognition. Considering that a ketogenic diet and intermittent fasting often face challenges in lasting adherence (Włodarek, 2019), BHB supplementation may yield more satisfactory long-term outcomes to prevent both weight gain and cognitive impairments linked to metabolic diseases.

It is important to note that the ketogenic diet has been utilized for a century as a protective strategy in epileptic patients (Fadó et al., 2022). However, this has been attributed to the medium-chain triglyceride decanoic acid, which is often elevated in this type of diet and blocks seizures through the direct inhibition of AMPARs, rather than ketone bodies themselves (Augustin et al., 2018).

In summary, our findings demonstrate that BHB effectively mitigates the adverse effects of PA on synaptic GluA1 abundance, excitability, and plasticity. Additionally, BHB administration in mice successfully prevents the cognitive decline associated with saturated high-fat, moderate-carbohydrate diet, which closely resembles the Western diet commonly consumed worldwide (characterized by a high intake of animal-based foods, fats, and simple sugars). Therefore, BHB supplementation could be a promising nutritional strategy to counteract the negative impacts of a Western diet on cognitive function.

## Materials and methods

### Animals and study design

All animal procedures performed at Universidad de Barcelona, by the Local Ethical Committee (#222/18) in agreement with European guidelines (2010/63/EU), while animal procedures at the Albert Einstein College of Medicine were approved by the Institutional Animal Care and Use Committee in accordance with the National Institutes of Health guidelines. C57BL/6J mice were housed in a controlled environment with a 12-hour light- 12-hour dark cycle, maintaining consistent temperature and humidity. At the age of 5 weeks, mice were randomly divided into two groups: one was fed a CD (10% of kcal from fat and 70% from carbohydrates; D12450J, Research Diets), while the other received a SFAD (49% of kcal from fat and 31% from carbohydrates; D19121204, Research Diets). Moreover, each group was subdivided into two subgroups: one received intragastrical administration of BHB (H6501, Sigma; 100 mg/kg/day), and the other received an equivalent volume of the vehicle (water). Throughout the study, animals had unrestricted access to food and water. After 7 weeks, the behavior tests were conducted, and mice were euthanized by decapitation 3 days later. Tissue samples were collected and stored at -80°C for subsequent analysis.

### Primary neuronal cultures

Primary cortical or hippocampal neurons were derived from mouse E16 embryos and obtained as described previously by (Parcerisas et al., 2020). Neurons were cultured in plates coated with poly-D-lysine (at 0.05 mg/ml for plastic or 0.1 mg/mL for coverslips; P7886, Sigma) and maintained in Neurobasal medium (21103049, Gibco), supplemented with B27 (17504044, Gibco), glutaMAX (35050061, Gibco), and antibiotics. Cultured neurons were maintained under controlled conditions at 37°C, 5% CO_2_, and 95% humidity. Every 3-4 days, one-fifth of the medium was replaced.

### Cellular treatments

Stock solution of 3mM OA (O7501, Sigma), PA (P9767, Sigma), and DHA (D8768, Sigma) were prepared at in defatted BSA solution (0.1 g/mL in 0.9% NaCl), as described previously (Fosch et al., 2023). Additionally, 250 nM BHB was prepared in water.

### Immunostaining, image acquisition, and analysis

IC was conducted on fixed cells using 4% (wt/vol) paraformaldehyde (252549, Sigma) as previously described (Casas et al., 2020). For surface GluA1 immunostaining, non-permeabilized cells were directly blocked with goat serum (G9023, Sigma), followed by the primary antibody incubation. For PSD95, cells were permeabilized using 0.1% Triton X-100 (vol/vol) before blocking and subsequent incubation with the primary antibody. Both mouse anti-GluA1 (1:300; ab174785) and rabbit anti-PSD95 (1:30; ab18258) were obtained from Abcam. Subsequent incubation was performed with the secondary antibody anti-rabbit Alexa488 (1:300; A11008) and anti-mouse Alexa568 (1:150; A11011), both purchased from Invitrogen. A brief 5-minute incubation with Hoechst solution (1 µg/mL; 14530, Sigma) was performed to visualize cellular nuclei. Finally, coverslips were mounted using Fluoromount Mounting Medium (F4680, Sigma).

Confocal imaging was conducted on immunostained cortical neurons using the Leica DMi8 confocal laser-scanning microscope with a ×63 1.4 NA oil objective. Cells were cultured, stained, and imaged simultaneously using identical settings for quantification. All subsequent quantifications were performed as previously described (Casas et al., 2020). The quantification of confocal images was performed on reconstructed 3D images using Imaris 9.2 software (Bitplane). To quantify surface GluA1 levels, primary dendrites, representing the initial branches extending from the cell soma, were selected. Then, a random region of interest (ROI) measuring 15 µm × 5 µm was chosen, and the integrated intensity was measured using the Spot tool in Imaris. To quantify synaptic GluA1 levels, the intensity of GluA1 staining within the 3D ROI defined by PSD95 (used as a synaptic marker) was quantified using the Surface tool in Imaris.

### AMPA-mediated miniature excitatory postsynaptic currents

mEPSCs were conducted in cultured cortical neurons seeded in coverslips, which were mounted in a recording chamber of an inverted microscope (Axio-Vert.A1 Zeiss). Whole– cell patch–clamp currents were recorded at room temperature (25-26°C) using an Axopatch 200B amplifier–Digidata1440A Series interface board with pClamp10 software (Molecular Devices) as described previously (Fadó et al., 2015). The extracellular solution used for bathing the cells contained the following (in mM): 140 NaCl, 3.5 KCl, 10 HEPES, 20 glucose, 1.8 CaCl2, and 0.8 MgCl2 (pH 7.4 adjusted with NaOH). To isolate AMPAR-mediated mEPSCs, specific blockers were added to the extracellular solution: 1 μM tetrodotoxin (ab120054, Abcam) to inhibit evoked synaptic transmission, 50 μM D-(-)-2-amino-5-phosphonopentanoic acid (ab120003 Abcam) to block NMDA receptor and 100 μM picrotoxin (P1675, Sigma) to block GABAA receptors. The intracellular solution contained (in mM): 116 K-glutonate, 6 KCl, 8 NaCl, 10 HEPES, 0.2 EGTA, 2 Mg-ATP, 0.3 Na-ATP (pH 7.2 adjusted with KOH). Series resistance was typically 17.71 ± 0.92 MΩ and was controlled at the experiment’s beginning and end. Cells exhibiting a change in resistance greater than 20% were excluded from the analysis. Data was digitized at 5 KHz and filtered at 2 KHz. mEPSCs were detected using an amplitude threshold of 5 pA. Only events with monotonic fast rise and uncontaminated decay were included for the amplitude. mEPSCs were analyzed with IgorPro 6.06 (WaveMetrics) using NeuroMatic 2.03 (Rothman & Silver, 2018).

### Hippocampal slice preparation and electrophysiology

Acute slice preparation and electrophysiology recording were conducted as described previously (Monday et al., 2022). One-month-old male mice were deeply anesthetized with isoflurane and then decapitated. The brain was rapidly removed from the skull and submerged in a sucrose-based solution. Sucrose-based solution contained the following composition (in mM): 215 sucrose, 2.5 KCl, 26 NaHCO3, 1.6 NaH2PO4, 1 CaCl2, 4 MgCl2, 4 MgSO4, and 20 D-glucose.Both hippocampi were carefully dissected. Transverse hippocampal slices (300 μm thick) were obtained by using a vibratome (VT1200s, Leica Microsystems). Then, slices were obtained and maintained at 34 °C for 30 minutes in artificial cerebrospinal fluid (aCSF) and at room temperature for at least 60 min before experimental procedures. aCSF, continuously bubbled with carbogen (95% O2 /5% CO2 mixture), contained the following (in mM): 124 NaCl, 2.5 KCl, 26 NaHCO3, 1 NaH2PO4, 2.5 CaCl2, 1.3 MgSO4, and 10 D-glucose.

Hippocampal slices were placed in an immersion chamber perfused with aCSF (2 mL/min) and supplemented with 50 μM picrotoxin to block GABA_A_-mediated inhibition. All experiments were performed at 28 ± 1°C. For fEPSP, a stimulating patch-type pipettes filled with aCSF and a recording patch-type pipette filled with 1M NaCl were placed in the *stratum radiatum* of CA1 with a distance of ∼100 μm between them. Synaptic responses were elicited by square-wave voltage or current pulses (100 μs pulse width) delivered through a stimulus isolator (Isoflex, AMPI) at 0.05 Hz. To build I/O curves, stimulus from 10 to 90 volts were used. The PPR was calculated as the ratio between the slopes of the second response by the first one. LTP was induced using a high-frequency stimulation protocol consisting of two trains of 100 pulses, 100 Hz delivered with a 10-second interval between them. Whole-cell patch-clamp recordings were performed in CA1 pyramidal cells. The patch pipettes were pulled from borosilicate glass (tip resistance 4-7 MΩ) and recordings were performed using a Cs^+^-based internal solution. Cs+-based internal solution contained the following (in mM): 131 Cs-gluconate, 8 NaCl, 1 CaCl2, 10 EGTA, 10 D-glucose, and 10 HEPES, at pH 7.2 (285-290 mOsm). Series resistance (15–25 MΩ) was monitored throughout all experiments with a -5 mV, 80 ms voltage step, and cells that exhibited a significant change (>20%) were excluded from the analysis. Excitatory postsynaptic currents (EPSCs) were evoked with an extracellular electrode placed in the *stratum radiatum* of CA1. The AMPAR-mediated EPSCs were recorded at −70 mV. After 5-10 minutes of recording, the cell was switched to +30 mV to record NMDA receptor-mediated EPSCs. The AMPA/NMDA ratio was calculated by dividing the peak amplitude of AMPA EPSC by NMDA EPSC. Previously, electronic subtraction was performed to remove any residual AMPA component from the recordings conducted at +30 mV. All recordings were performed using a Multi-clamp 700B amplifier (Molecular Devices). Data were acquired at 5 kHz, filtered at 2.4 kHz and analyzed offline using IgorPro 7 (WaveMetrics).

### BHB determination

According to the manufacturer’s protocol, BHB levels were measured using a BHB Assay kit (MAK041, Sigma). The amount of BHB present was then normalized by the tissue weight.

### Behavior tests

NORT was used to evaluate short (2h) and long-term memory (24h) as described previously by (Griñán-Ferré et al., 2020). For NORT a 90o, two-arm, 25 cm-long, and 20 cm-high maze was used. Before the test, mice were individually submitted to a 3-day habituation period in the maze, 10 minutes daily. On day 4, they were allowed to explore the maze during a 10-minute acquisition trial (habituation phase), examining two identical novel objects (O1 and O1’) placed at the end of each arm. 2 hours later, short-term memory was evaluated with another 10-minute retention trial, where one of the two identical objects was replaced by a novel one (TO). 24 hours later, long-term memory was tested again, using a new object (TN) and one used in the previous trial (TO). The exploration time of the novel object (TN) and the exploration time of the old object (TO) were quantified by visualizing video recordings of each trial session. Object exploration was defined as the mouse’s nose touching the object or pointing it toward the object at a distance ≤ 2cm. The cognitive performance was analyzed using the DI calculated as (TN-TO)/(TN+TO).

OLT was employed to evaluate hippocampus-dependent spatial learning using a box (50 x 50 x 25 cm) with three white and one black walls. The box remained empty the first day, allowing mice to acclimate for 10 minutes. The following day, two identical 10-cm-high objects were positioned equidistant from each other and the black wall for 10 minutes. On the last day, one of the objects was relocated in front of the opposite white wall to test spatial memory, quantified as with NORT.

OFT was used to analyze locomotor activity and anxiety-related behaviors as described previously (Griñan-Ferré et al., 2016). For OFT, the box (50 x 50 x 25 cm) was divided into central and peripheral zones to facilitate the analysis of the distance traveled. Mice were positioned either at the center or at one of the corners of the open field. After 5 minutes of exploration time, the distance traveled and the number of rearings were analyzed.

### Statistical analysis

All results are shown as the mean ± standard error (SEM) or standard deviation (SD). Statistical analyses were conducted using the GraphPad Prism 9.0 Software. According to the normality test (Shapiro-Wilk test and D’Agostino and Pearson test), Student’s t-test or Mann-Whitney U test were used when comparing only two groups. If multiple groups with different variables were compared, One-way or Two-way analysis of variance (ANOVA) was performed, followed by Bonferroni tests to assess specific group differences.

## Acknowledgements

This study was supported by the Spanish Ministry of Science and Innovation (MCIN) (PID2020-114953RB-C22 to N.C.; PID2022-139016OA-I00 and PDC2022-133441-I00 to C.G-F. and M.P.; PID2020-11751ORB-I00 to J.R.A.; PID2020-119932GB-I00 and María de Maeztu Unit of Excellence CEX2021-001159-M to M.P. and D.S.), the CIBEROBN (CB06/03/0001 to N.C.), the FEDER Regional European Development Fund and CIBERNED (CB06/05/0042 to J.R.A.), the Generalitat de Catalunya (2021 SGR 00357 to C.G-F. and M.P.), and the NIH grants (R01MH116673, R01MH125772, and R01NS113600 to P.E.C. and E.G.).

## Author Contributions

Rocío Rojas: Investigation, Formal analysis, Data curation, Conceptualization. Writing – original draft. Christian Griñán-Ferré: Investigation, Funding acquisition. Aida Castellanos: Investigation. Ernesto Griego: Investigation. Marc Martínez: Investigation. Juan de Dios Navarro-López, Lydia Jiménez-Díaz, José Rodríguez-Álvarez, and Mercè Pallàs: Funding acquisition. David Soto del Cerro: Investigation, Formal analysis, Funding acquisition. Pablo E Castillo: Funding acquisition, Supervision. Rut Fadó: Investigation, Formal analysis, Data curation, Conceptualization, Supervision, Writing – original draft. Núria Casals: Conceptualization, Funding acquisition, Project administration, Supervision, Writing – review & editing.

## Additional files

- ARRIVE guidelines 2.0: Essential 10 checklist
- Supplemental figures from 1 to 5
- Supplemental materials and methods

