## Supplemental materials and methods for "BETA-HYDROXYBUTYRATE COUNTERACTS THE DELETERIOUS EFFECTS OF A SATURATED HIGH-FAT DIET ON SYNAPTIC AMPA RECEPTORS AND COGNITIVE PERFORMANCE"

### Cell line and treatment

SH-SY5Y cells, derived from SH-SY subclone of the parental SK-N-SH human neuroblastoma cell line, were cultured in Dulbecco's Modified Eagle's Medium (DMEM; D5671, Sigma) supplemented with 10% fetal bovine serum (FBS12A, Capricorn Scientific), 1% glutamax (X0551-100, Biowest), 100 U/ mL penicillin and 100 µg/mL streptomycin (P0781, Sigma), and maintained at 37°C in 5% CO2 humidified air. Cell line was tested for mycoplasma contamination using EZ-PCR™ Mycoplasma Detection Kit (20-700-20BI). To induce neuron-like phenotype, cells were seeded at 40,000 cells/mL in collagen-coated coverslips (1:50; 354236, Corning) and cultured in medium supplemented with 10 µM of retinoic acid (R2622, Sigma) diluted in DMSO (1029521011, Merck). Differentiated SH-SY5Y cells were subjected to treatment 6 days after, with the specific final concentrations as indicated in results section.
