## Supplemental figures for "BETA-HYDROXYBUTYRATE COUNTERACTS THE DELETERIOUS EFFECTS OF A SATURATED HIGH-FAT DIET ON SYNAPTIC AMPA RECEPTORS AND COGNITIVE PERFORMANCE"

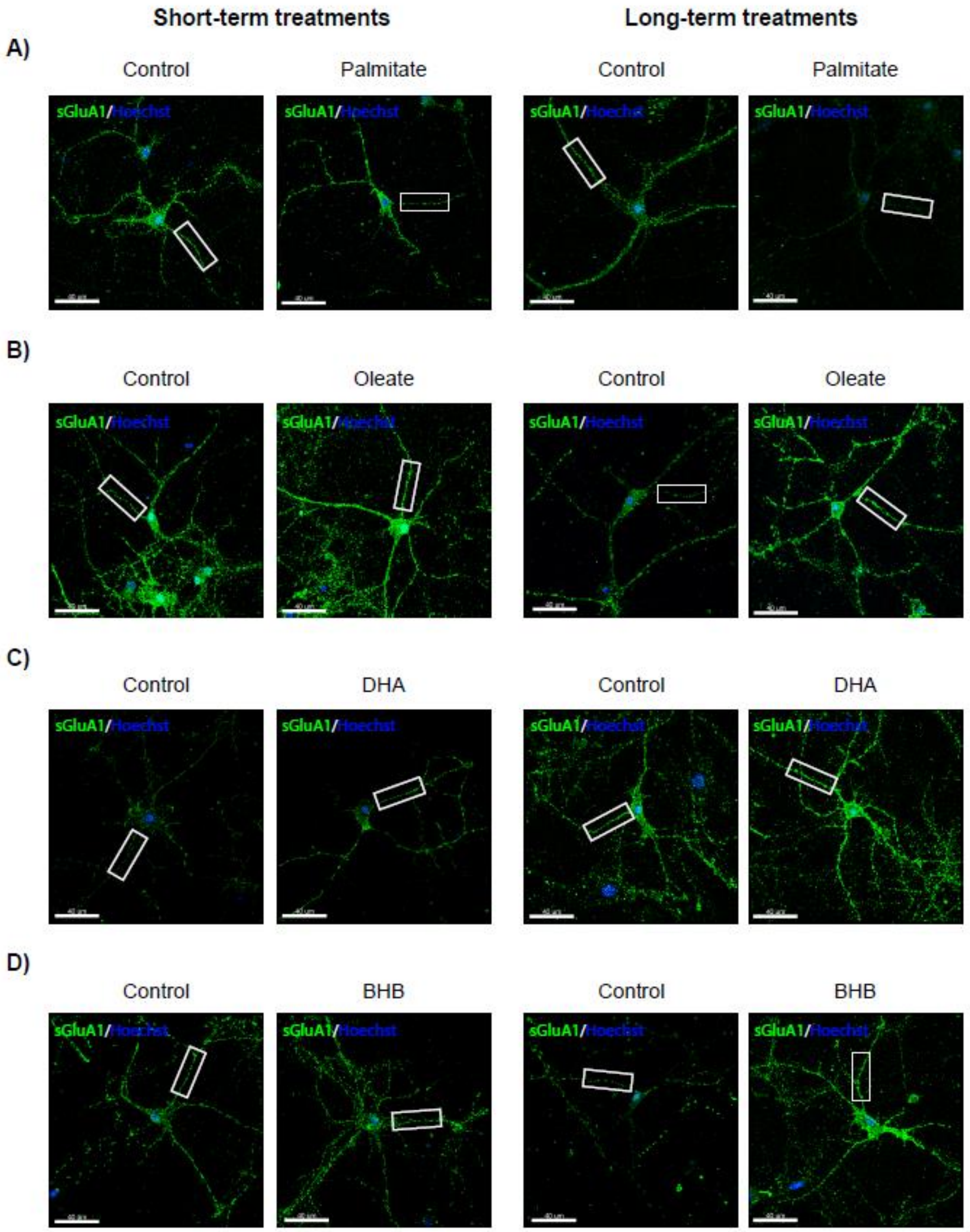


**Supplemental Figure 1.** Complete images of nutrient's effects on surface GluA1 levels in primary cortical neurons. Cortical neurons at 13-14 DIV were submitted to PA (200 µM; **A, B**), OA (200 µM; **C, D**), DHA (200 µM; **E, F**), or BHB (5mM; **G, H**) treatment for 2 or 24 hours, respectively. Surface GluA1 was detected by IC (green stain) and nuclei by Hoechst staining (in blue). The magnifications of images are shown in Figure 1. Scale bar = 40 µm

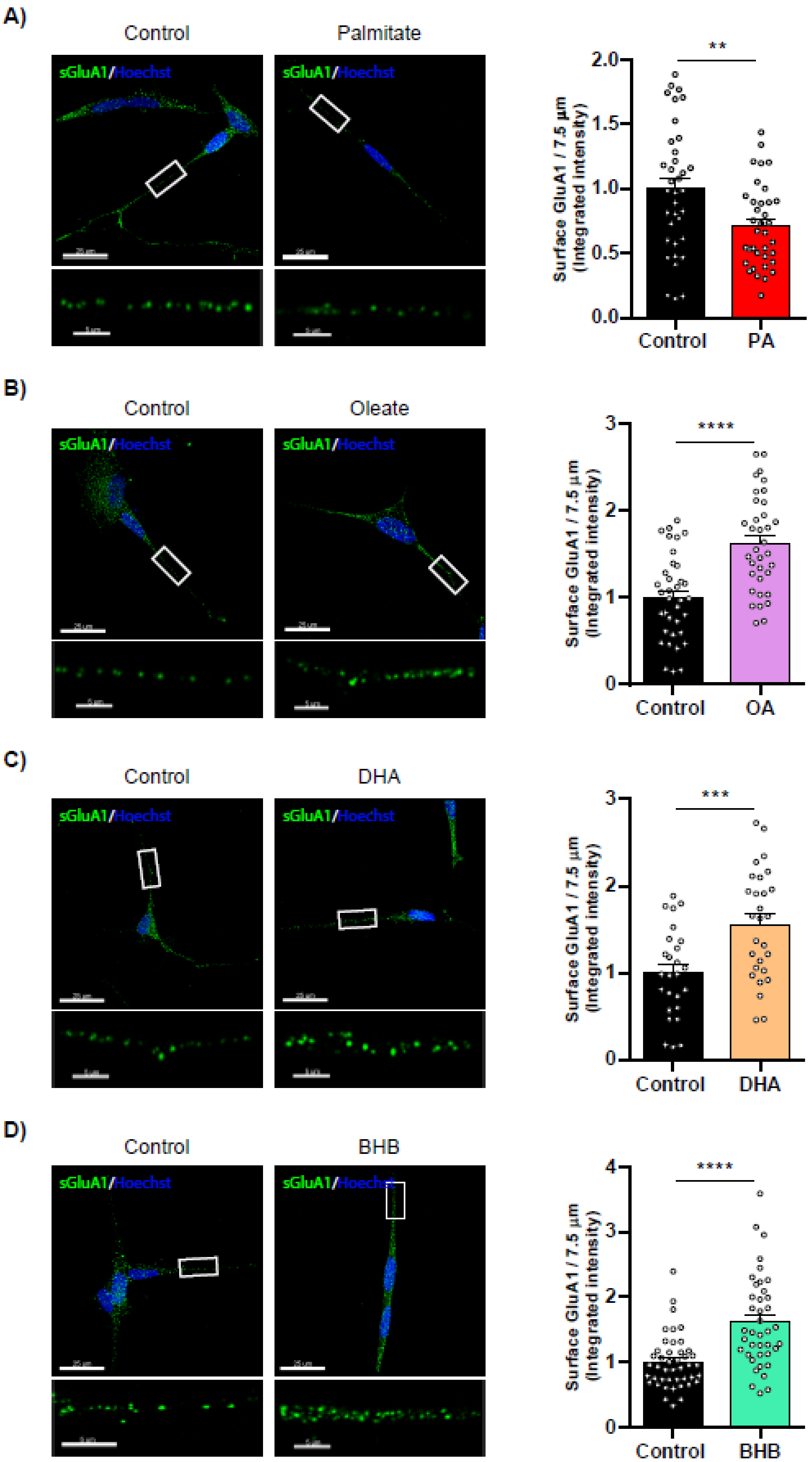
A)
Control
Palmitate
sGluA1/Hoechst
sGluA1/Hoechst
Surface GluA1 / 7.5 µm (Integrated intensity)
2.0
1.5
1.0
0.5
0.0
**
Control
PA
B)
Control
Oleate
sGluA1/Hoechst
sGluA1/Hoechst
Surface GluA1 / 7.5 µm (Integrated intensity)
3
2
1
0
****
Control
OA
C)
Control
DHA
sGluA1/Hoechst
sGluA1/Hoechst
Surface GluA1 / 7.5 µm (Integrated intensity)
3
2
1
0
***
Control
DHA
D)
Control
BHB
sGluA1/Hoechst
sGluA1/Hoechst
Surface GluA1 / 7.5 µm (Integrated intensity)
4
3
2
1
0
****
Control
BHB

**Supplemental Figure 2.** Differentiated SH-SY5Y successfully mimics nutrients' effects on surface GluA1 levels. The SH-SY5H cell line was differentiated to a neuron-like phenotype with retinoic acid (10µM). 6 days later, cells were exposed for 24 hours to PA (200 µM; **A**), OA (200 µM; **B**), DHA (200 µM; **C**), or BHB (5mM; **D**), respectively. Then, nonpermeabilized cells were fixed and processed, as in Figure 1. Representative images of neurites are shown on the right. In each condition, we analyzed 22-35 neurites from one (DHA) or two (PA, OA, and BHB) independent experiments performed by biological duplicates. Results in left graphs are given as mean ± SEM. (**A**) Control (BSA; 1.00 ± 0.08, n = 36), and PA (0.71 ± 0.05, n = 36). (**B**) Control (BSA; 1.00 ± 0.08, n = 36), and OA (1.63 ± 0.09, n = 35). (**C**) Control (BSA; 1.00 ± 0.10, n = 26), and DHA (1.56 ± 0.12, n = 28). (**D**) Control (1.00 ± 0.06, n = 44), and BHB (1.62 ± 0.11, n = 40). **, P < 0.05; ***, P < 0.001; ****, P < 0.0001. Student t-test. Scale bar = 25 µm; scale bar of inset magnifications = 5 µm. *p<0.05.

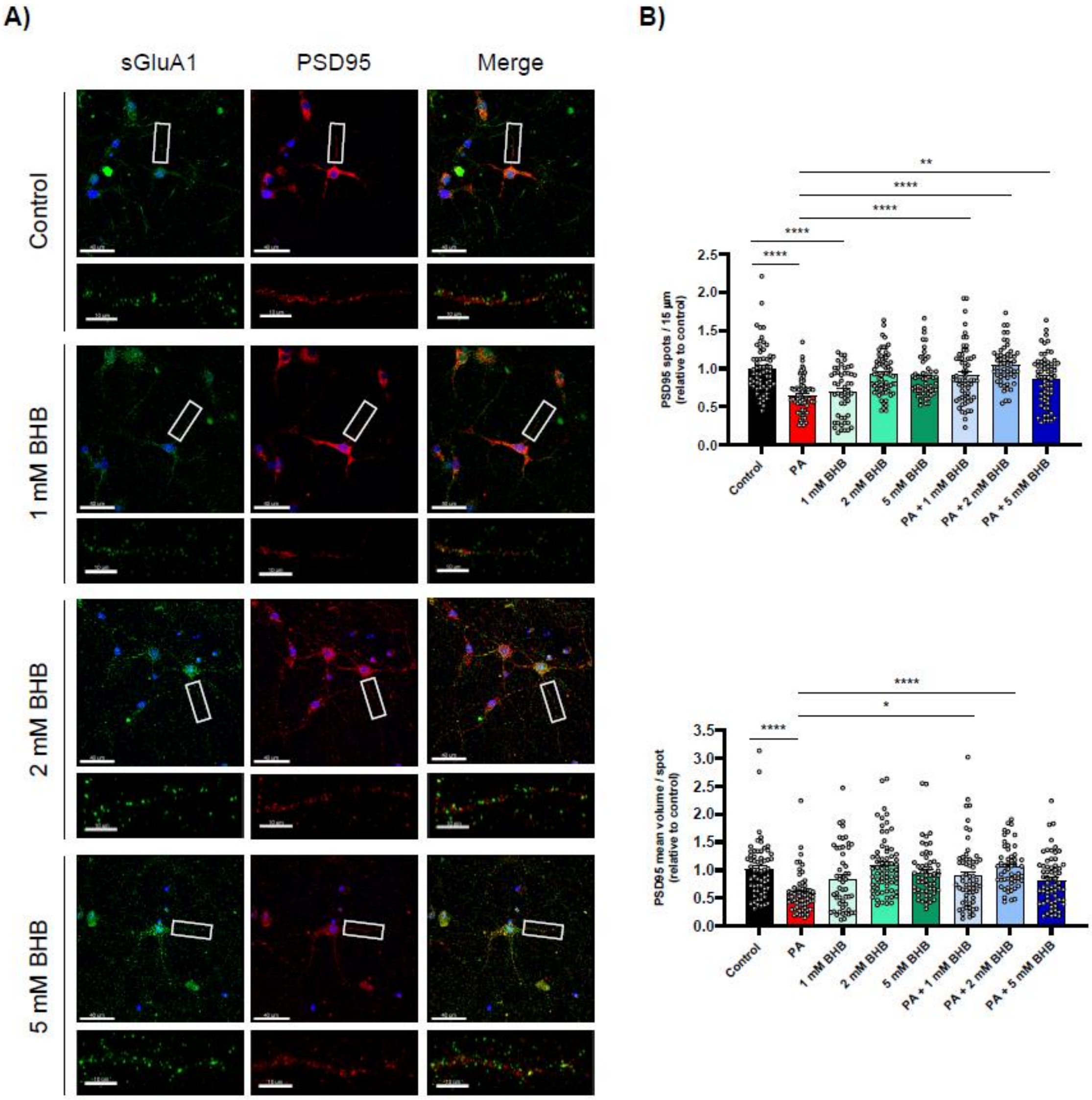


**Supplemental Figure 3. PA decreases, and high-dose BHB increases surface**

**GluA1 levels in hippocampal neurons.** (A) Representative images from the same experiments done in Figure 3 are shown. (B) PSD95 number and volume of spots. Results in the graphs are given as mean ± SEM of 60 dendrites from two independent experiments performed by biological duplicates. PSD95 spots (upper graph): Control (BSA; 1.00 ± 0.04, n=61), PA (0.65 ± 0.03, n=63), 1 mM BHB (0.69 ± 0.05, n=51), 2 mM BHB (0.92 ± 0.03, n=68), 5 mM BHB (0.88 ± 0.04, n=56), PA + 1 mM BHB (0.92 ± 0.05, n=61), PA + 2 mM BHB (1.05 ± 0.04, n=53), and PA + 5 mM BHB (0.86 ± 0.04, n=63). PSD95 volume (below graph): Control (BSA; 1.00 ± 0.07, n=61), PA (0.58 ± 0.04, n=63), 1 mM BHB (0.85 ± 0.08, n=51), 2 mM BHB (1.08 ± 0.06, n=68), 5 mM BHB (0.96 ± 0.06, n=56), PA + 1 mM BHB (0.90 ± 0.07, n=61), PA + 2 mM BHB (1.06 ± 0.05, n=53), and PA + 5 mM BHB (0.82 ± 0.06, n=63). Scale bar = 40 µm; scale bar of inset magnifications = 10 µm. ****, P < 0.0001. One-way ANOVA followed by Bonferroni test.

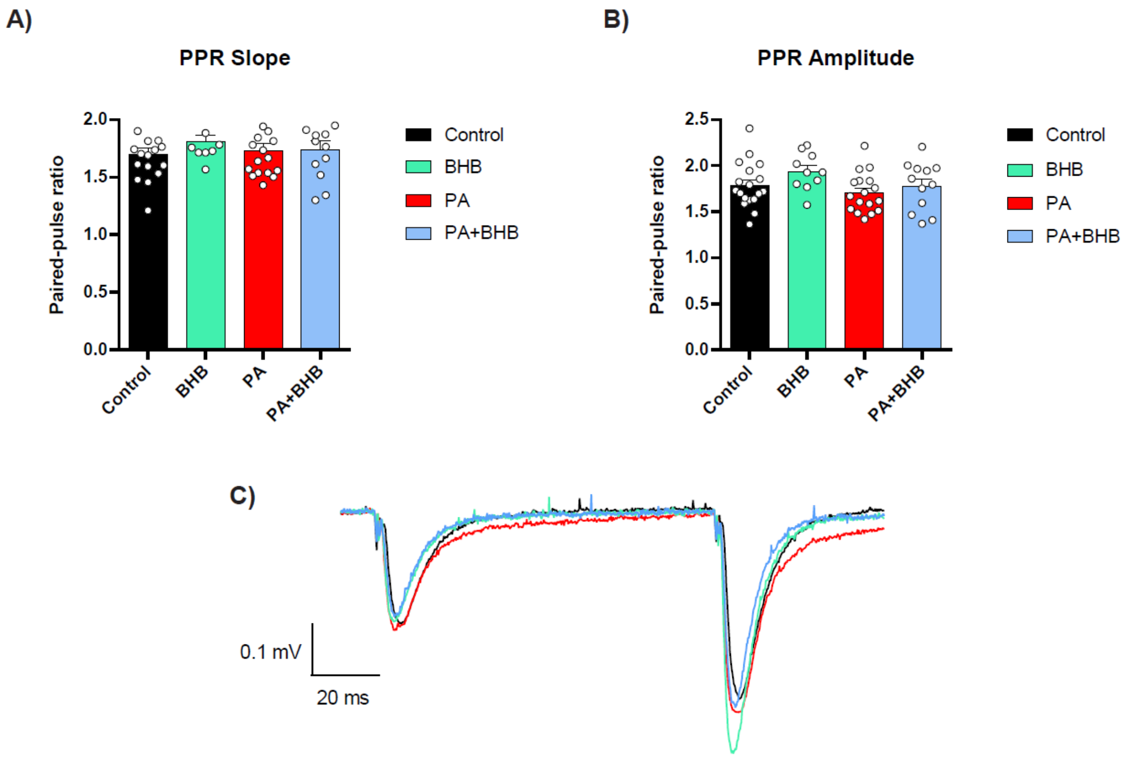


**Supplemental Figure 4. Effects of BHB, PA, and co-treatment of hippocampal slices on paired-pulse ratio.** Average values of PPRs from fEPSP with 100 ms interstimulus interval. Individual values for each condition are displayed as a dot plot. Data are represented as mean ± SEM of n = 10-17 slices from 4 male mice (P26 ± 5) per condition. Representative traces for each condition are shown. PPR slope: control (1.70 ± 0.05, n=17), BHB (1.81 ± 0.06; n=9), vehicle (1.78 ± 0.07, n=12), PA (1.73 ± 0.06, n=17), and PA + BHB (1.74 ± 0.08, n=12). PPR amplitude: control (1.79 ± 0.06, n=18), BHB (1.94 ± 0.06; n=10), vehicle (1.79 ± 0.06, n=12), PA (1.71 ± 0.05, n=17), and PA + BHB (1.78 ± 0.08, n=12). One-way ANOVA followed by Bonferroni test.

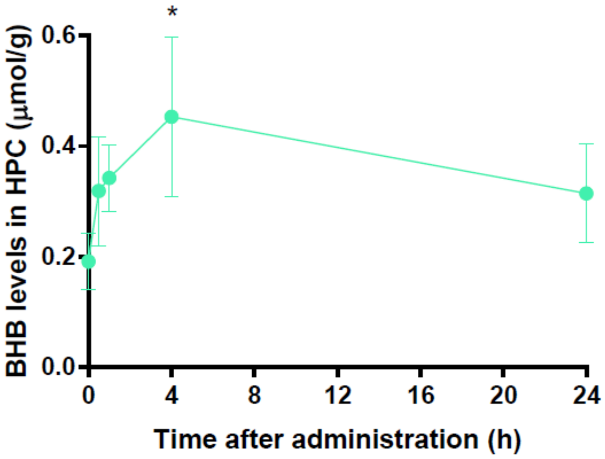


**Supplemental Figure 5. BHB hippocampal levels after BHB administration.** BHB pharmacokinetic assay. BHB levels were measured in the hippocampus of mice at 30 minutes, 1 hour, 2 hours, 4 hours, and 24 hours following intragastric administration of BHB (100 mg/kg). Results are represented as mean ± SD from n=2-4/time point. *, $P < 0.05$. One-way ANOVA followed by Bonferroni test.
